# Lipid droplets fuel SARS-CoV-2 replication and production of inflammatory mediators

**DOI:** 10.1101/2020.08.22.262733

**Authors:** Suelen da Silva Gomes Dias, Vinicius Cardoso Soares, André C. Ferreira, Carolina Q. Sacramento, Natalia Fintelman-Rodrigues, Jairo R. Temerozo, Lívia Teixeira, Ester Barreto, Mayara Mattos, Caroline S. de Freitas, Isaclaudia G. Azevedo-Quintanilha, Pedro Paulo A. Manso, Eugenio D. Hottz, Camila R. R. Pão, Dumith C. Bou-Habib, Fernando A. Bozza, Thiago M. L. Souza, Patrícia T. Bozza

## Abstract

Viruses are obligate intracellular parasites that make use of the host metabolic machineries to meet their biosynthetic needs, identifying the host pathways essential for the virus replication may lead to potential targets for therapeutic intervention. The mechanisms and pathways explored by SARS-CoV-2 to support its replication within host cells are not fully known. Lipid droplets (LD) are organelles with major functions in lipid metabolism and energy homeostasis, and have multiple roles in infections and inflammation. Here we described that monocytes from COVID-19 patients have an increased LD accumulation compared to SARS-CoV-2 negative donors. *In vitro*, SARS-CoV-2 infection modulates pathways of lipid synthesis and uptake, including CD36, SREBP-1, PPARγ and DGAT-1 in monocytes and triggered LD formation in different human cells. LDs were found in close apposition with SARS-CoV-2 proteins and double-stranded (ds)-RNA in infected cells. Pharmacological modulation of LD formation by inhibition of DGAT-1 with A922500 significantly inhibited SARS-CoV-2 replication as well as reduced production of pro-inflammatory mediators. Taken together, we demonstrate the essential role of lipid metabolic reprograming and LD formation in SARS-CoV-2 replication and pathogenesis, opening new opportunities for therapeutic strategies to COVID-19.

## Introduction

The coronavirus disease 2019 (COVID-19) caused by the novel severe acute respiratory syndrome-coronavirus 2 (SARS-CoV-2) has rapidly spread in a pandemic, representing an unprecedented health, social and economic threat worldwide (Lu et al., 2020; Wu et al., 2020). This newly emerged SARS-CoV-2 belongs to the *Betacoronavirus* genus of the subfamily *Orthocoronavirinae* in the *Coronaviridae* family. Like other Coronavirus, the SARS-CoV-2 is an enveloped non-segmented positive-sense RNA (+RNA) virus (Zhu et al., 2020), which genome sequence is similar to the already known SARS-CoV (Zhou et al., 2020b). Despite the similarity with other members of the *Betacoronavirus* genus, the pathogenesis of SARS-CoV-2 infection presents unique properties that contribute to its severity and pandemic-scale spread. Therefore, it is necessary to understand how the virus interacts and manipulate host cell metabolism to develop novel strategies to control the clinical progression of the infection and to limit the SARS-CoV-2 pandemic.

Viruses are obligated intracellular pathogens that require host cell machinery to replicate (Chazal and Gerlier, 2003; Novoa et al., 2005; Takahashi and Suzuki, 2011). Viruses interact with several intracellular structures and have the ability to reprogram the cellular metabolism to benefit viral replication (Abrantes et al., 2012; Syed et al., 2010; Zhang et al., 2017). Accumulated evidence points to major roles of lipid droplets (LD) for virus replication cycle and pathogenesis, highlighting the potential of these organelles as targets for drug development. Many studies have demonstrated the interaction of viral molecules with LD-related components, and the relevance of these organelles for viral replication as already demonstrated in several positive-strand RNA (+ RNA) viruses such as *Flaviviridae* members, rotavirus and reovirus (Cheung et al., 2010; Coffey et al., 2006; Filipe and McLauchlan, 2015; Lyn et al., 2013; Samsa et al., 2009; Villareal et al., 2015). Accordingly, pharmacological interventions that alter the synthesis of fatty acids, enzymes associated with lipid metabolism and the formation of LDs reduce viral replication and assembly (Herker et al., 2010; Martín-Acebes et al., 2011; Villareal et al., 2015; Yang et al., 2008; Zhang et al., 2017). In addition, LDs play an important role in the infection pathogenesis and inflammatory processes (Herker and Ott, 2012; Pereira-Dutra et al., 2019).

Here we demonstrate major effects of SARS-CoV-2 to modulate cellular lipid metabolism in human cells favoring increased de novo lipid synthesis and lipid remodeling, leading to increased LD accumulation in human cells. We also reported increased LD accumulation in monocytes from COVID-19 patients when compared to healthy volunteers. Importantly, the blockade of LD biogenesis with a pharmacological inhibitor of DGAT-1 decreased viral replication and pro-inflammatory cytokine production and prevented cell death. Collectively, our results uncover mechanisms of viral manipulation of host cell lipid metabolism to allow SARS-CoV-2 replication and may provide new insights for antiviral therapies.

## Results

### SARS-CoV-2 infection upregulates lipid metabolism, increasing LD biogenesis in human cells

Viruses have the ability to modulate cellular metabolism with benefits for their own replication. Several +RNA viruses, including members of *Flaviviridae* family, as HCV (Boulant et al., 2007; Lyn et al., 2013) and DENV (Carvalho et al., 2012; Samsa et al., 2009), as well as reovirus (Coffey et al., 2006) and poliovirus (Viktorova et al., 2018) modify the lipid metabolism in different cells and trigger LD formation, using these host organelles at different steps of their replicative cycle.

Here, we demonstrated increased LD accumulation in human monocytes from COVID-19 patients when compared with healthy volunteers (Fig. 1A and B). Likewise, we demonstrated that in vitro infection with SARS-CoV-2 with a multiplicity of infection (MOI) of 0.01 triggers the increase of LDs in primary human monocytes within 24 hours (Fig. 1C and D), as well as in a human lung epithelial cell line (A549), and human lung microvascular endothelial cell line (HMVEC-L) after 48 hours post-infection (Supplementary Fig. 1).

**Fig 1.**
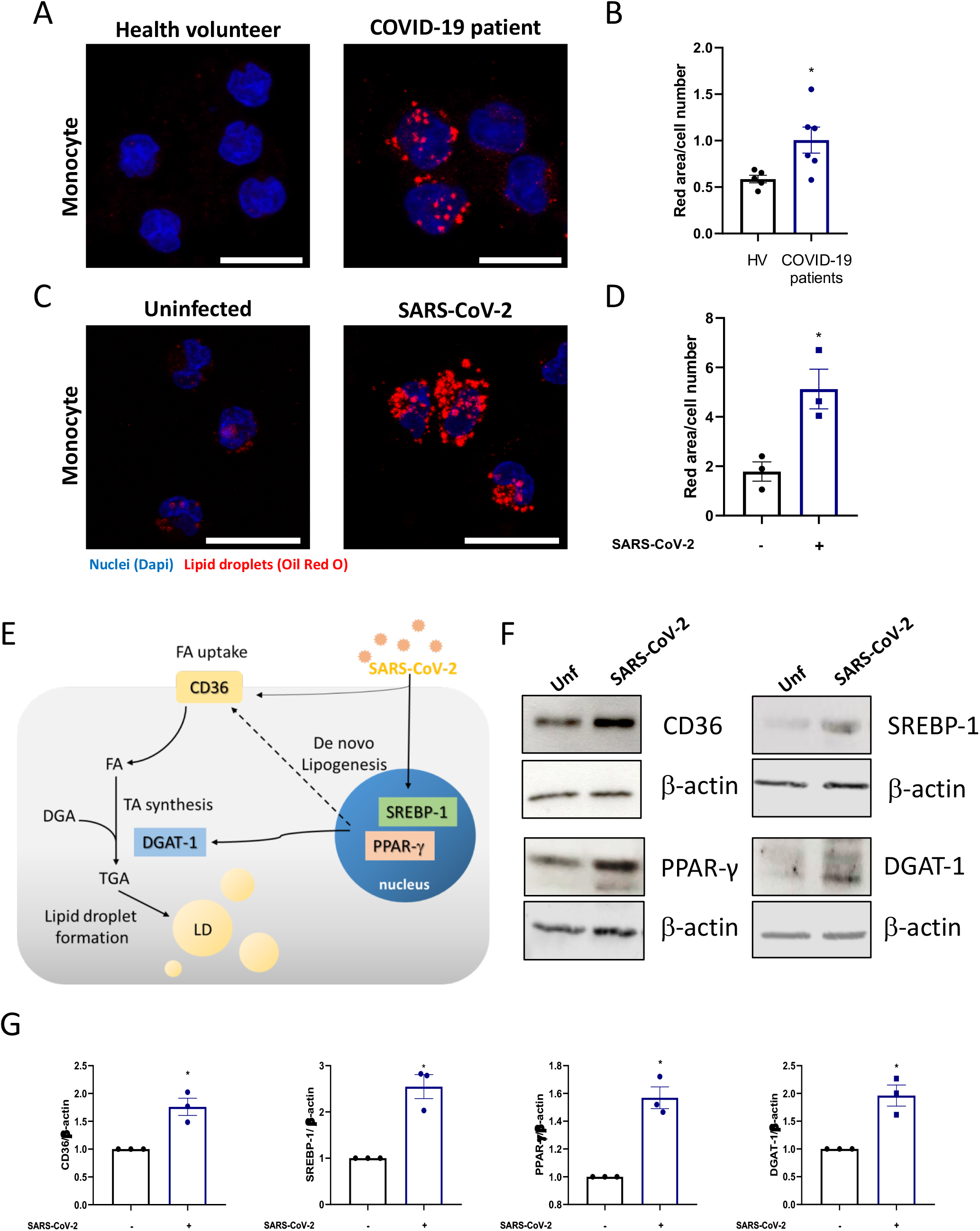
SARS-CoV-2 infection modulates the lipid metabolism in human monocytes. (A and C) LDs were captured by fluorescent microscopy after Oil Red O staining (Red) and nuclei stained with DAPI (Blue). (A) Representative images of monocytes from COVID-19 patients and health volunteers. (C) Representative images of human monocytes obtained from PBMC infected by SARS-CoV-2 with MOI of 0.01 for 24 hours. Scale bar 20μm. (B and D) LDs were evaluated by ImageJ software analysis by the measurement of the fluorescent area. (E) Representative scheme of the increase of proteins associated with lipid metabolism by SARS-CoV-2 infection in monocyte can regulate the lipid droplet formation. (F) Monocytes were infected by SARS-CoV-2 with MOI of 0.01 during 24h. Cell lysates were collected for the detection of CD36, PPAR-γ, SREBP-1, DGAT-1 by Western blotting. β-actin level were used for control of protein loading. (G) Densitometry data set of each protein. Data are expressed as mean ± SEM of five healthy volunteers (HV) for ex vivo experiments and three healthy donors for LDs staining and western blot. *p < 0.05 versus health volunteers or uninfected cells.

Lipid metabolism alterations in cells and plasma are emerging as major phenotypes during COVID-19 and SARS-CoV-2 infection (Shen et al., 2020). To gain insights on the mechanisms involved in LD formation, we evaluated whether SARS-CoV-2 infection could modulate the expression of the proteins associated with lipid metabolism involved in lipid uptake and de novo lipid synthesis (Fig. 1E). As shown in figure 1E-G, SARS-CoV-2 infection of human primary monocytes up-regulated the pathways involved in lipid uptake such as CD36, the major transcriptional factors involved in lipogenesis, PPARγ and SREBP-1, and the enzyme DGAT-1, which is involved in triacylglycerol synthesis, after 24 hours of infection.

Altogether, these data suggest that SARS-CoV-2 is able to modulate multiple pathways of lipid metabolism and remodeling, including in immune cells from COVID-19 patients, culminating in new LD assembling in human cells.

### Inhibition of LD formation decreases viral replication and prevents cell death in SARS-CoV-2 infected monocytes

DGAT-1 is a key enzyme involved in the final step of triacylglycerol synthesis and thus is central to remodel and finish the biogenesis of LDs (Chitraju et al., 2017). During HCV infection, DGAT-1 was shown to be required for LD biogenesis, and to control HCV protein trafficking to LDs (Camus et al., 2013). Consequently, DGAT-1 inhibition blocks HCV use of LDs as replication platforms and inhibits viral particle formation (Camus et al., 2013; Herker et al., 2010). To assess the involvement of DGAT-1 in LD biogenesis during the SARS-CoV-2 infection, we treated A549 cells with A922500, an inhibitor of the enzyme DGAT-1, for 2 hours at different concentrations prior to SARS-CoV-2 infection and evaluated the LD biogenesis 48 hours after infection. As shown in figure 2A and B, treatment with A922500 inhibited in a dose dependent manner the LD formation triggered by SARS-CoV-2 infection. Similarly, pre-treatment with A922500 also blocked LD induced by SARS-CoV-2 in monocytes (Fig. 2A and C).

**Fig 2.**
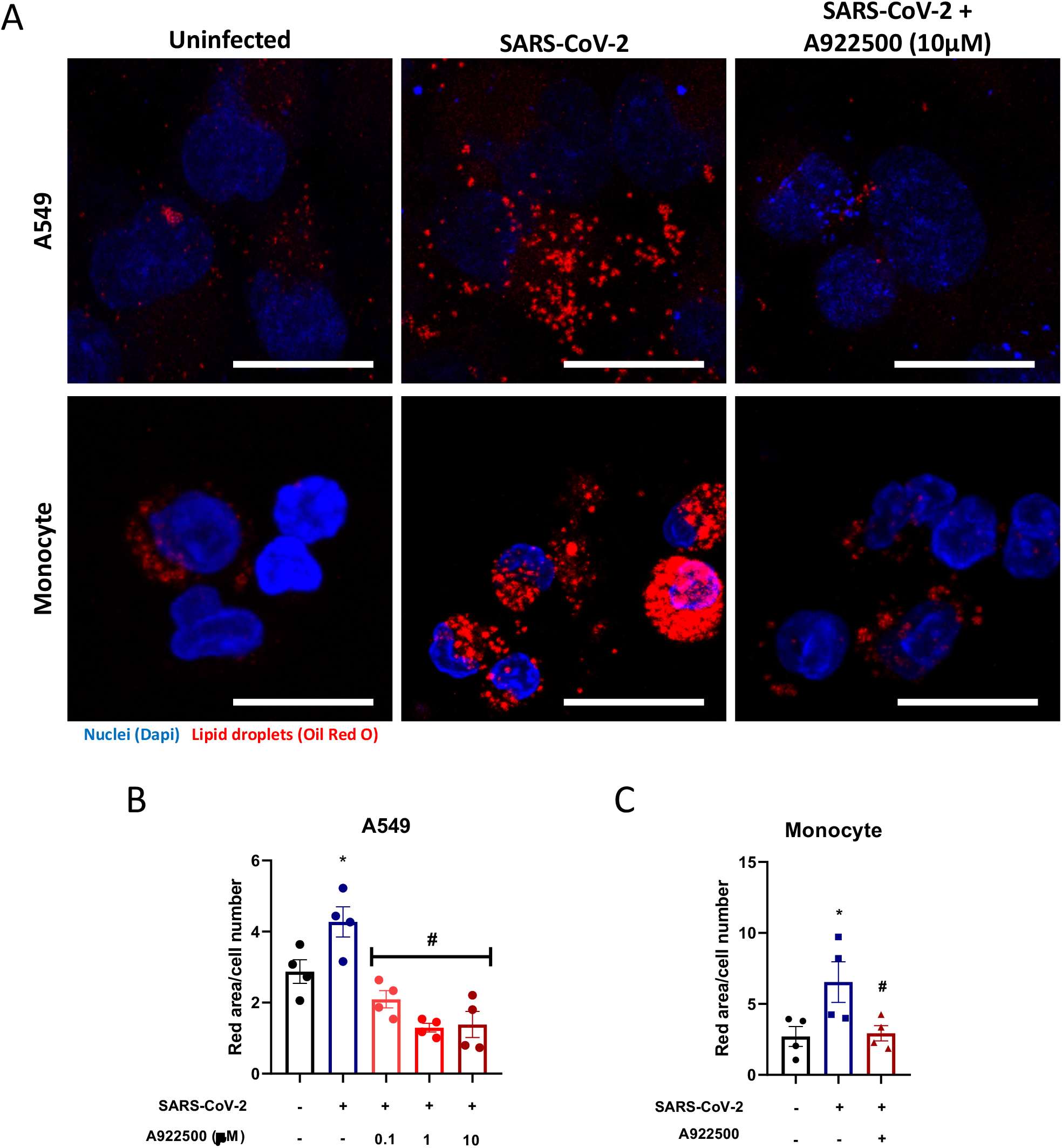
The A922500 inhibits lipid droplet biogenesis induced by SARS-CoV-2 in human pulmonary cells and monocytes. Human pulmonary cell (A549 cell line) and monocytes were pre-treated with DGAT-1 inhibitor A922500 for 2 hours before the infection with SARS-CoV-2 at MOI of 0.01 during 24h in monocytes and 48h in A549 cell line. (A) LDs were captured by fluorescent microscopy after Oil Red O staining (Red) and nuclei stained with DAPI (Blue). Scale bar 20μm. (B and C) LDs were evaluated by ImageJ software analysis by the measurement of the fluorescent area of (B) A549 pre-treated with A922500 using different concentrations (0.1, 1 and 10μM) and (C) LDs from monocytes pre-treated with A922500 (10μM). Data are expressed as mean ± SEM obtained in four independent experiments or donors. *p < 0.05 *versus* uninfected cells and #p < 0.05 *versus* A922500 treated cells.

Human monocytes infected with SARS-CoV-2 were shown to sustain viral genome replication, express higher levels of pro-inflammatory cytokines and may undergo cell death (Codo et al., 2020). To gain insights on the functions of LDs in SARS-CoV-2 infection, LD biogenesis was inhibited by A922500, a DGAT-1 inhibitor. Treatment with A922500 significantly reduced the viral load in human primary monocytes (Fig. 3A), suggesting a role for DGAT-1 and LD in SARS-CoV-2 replication.

**Fig 3.**
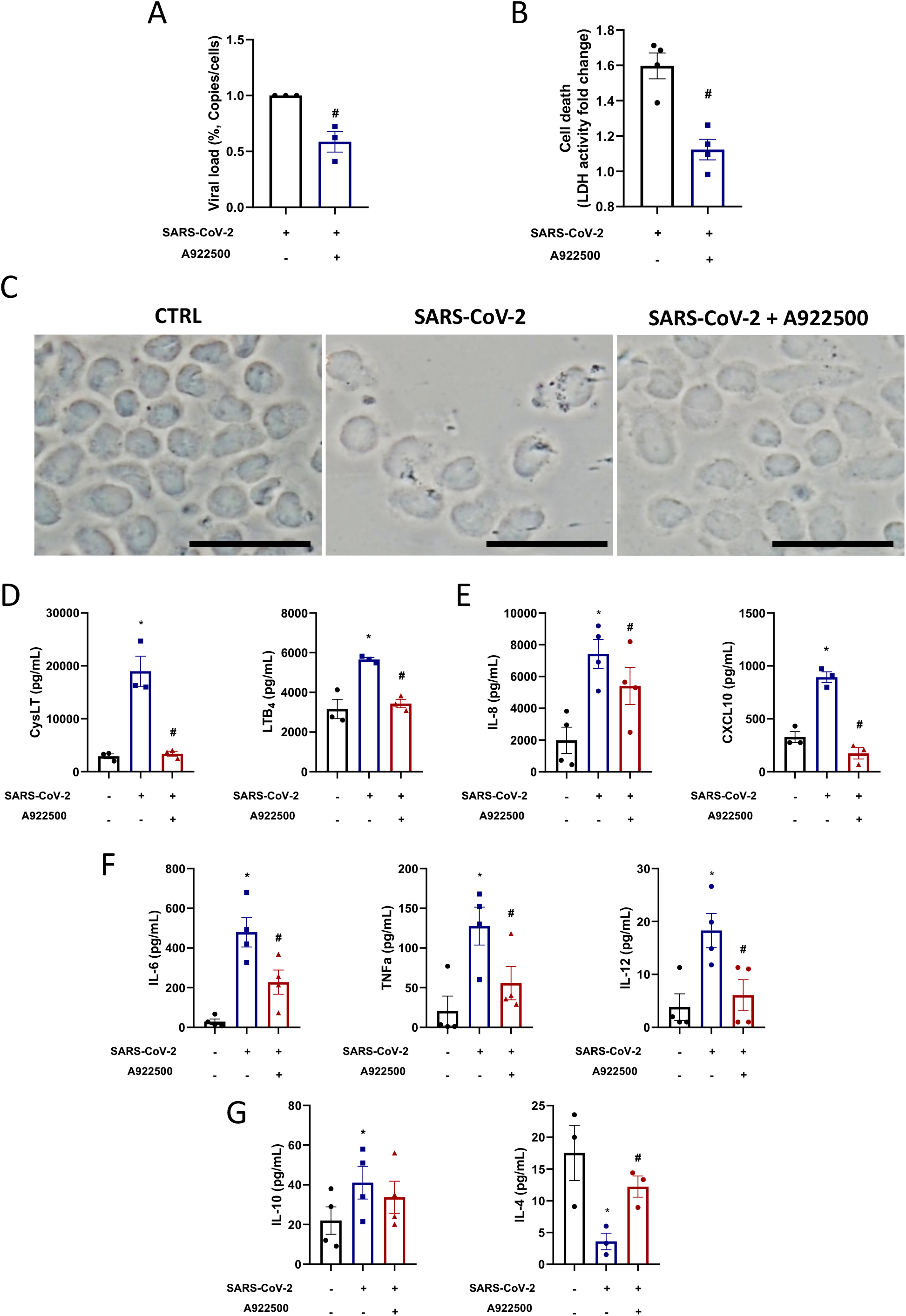
Inhibitor A922500 decreases the pro-inflammatory profile and cell death induced by SARS-CoV-2 infection and reduces the viral load in human monocyte. Monocytes were pre-treated with DGAT-1 inhibitor A922500 (10μM) for 2 hours before the infection with SARS-CoV-2 with MOI of 0.01 during 24h. (A) Cell death was measured in the supernatant by LDH activity fold change in relation to the uninfected cell. (B) Viral load by qPCR. Monocytes of each sample were counted for normalization. (C) Images of phase contrast from monocytes. Scale bar 20μm. (D-G) The inflammatory cytokines were measured in supernatants by ELISA (D) leukotrienes: CysLT and LTB_4_, (E) chemokines: IL-8 and CXCL-10, (F) pro-inflammatory: IL-6, TNF-α and IL-12 and (G) anti-inflammatory cytokines: IL-10 and IL-4. Data are expressed as mean ± SEM obtained in four independent donors. * p <0.05 *versus* uninfected cells and #p <0.05 *versus* A922500 treated cells.

It has already been demonstrated the capacity of the SARS-CoV-2 infection to induce cell death in monocytes, as evidenced by the leak of LDH to the extracellular space (Fintelman-Rodrigues et al., 2020). Cell death after viral infections can occur due to changes in cellular homeostasis caused by the virus replication per se and/or by the heightened inflammatory response. Here, we measured the death of human primary monocytes infected with SARS-CoV-2 by the release of LDH into the supernatant, and also by the analysis of cell morphology observed in phase contrast. Our data show that SARS-CoV-2 triggered increased LDH release in the supernatant and that infected cells exhibited morphologic alterations with membrane rupture/damage compatible with necrosis (Fig. 3B and C). Similar to the observed for viral replication, DGAT-1 and LD inhibition with 10 μM of A922500 was able to inhibit SARS-CoV-2-induced cell death (Fig. 3B and C).

### Lipid droplets are involved in SARS-CoV-2 heightened inflammatory response

Dysregulated immune response, with increased pro-inflammatory cytokine/chemokine production is observed during severe COVID-19 and associates with the outcome of the disease (Coperchini et al., 2020). We observed that primary human monocytes infected with SARS-CoV-2 exhibit increased production of leukotrienes (LTB_4_ and cysLT), pro-inflammatory cytokines (IL-6, TNFα and IL-12) and chemokines (IL-8 and CXCL10) in comparison with uninfected cells (Fig. 3D, E and F). Regarding anti-inflammatory mediators, SARS-CoV-2 infection increased IL-10 and reduced IL-4 production in comparison with uninfected monocytes (Fig. 3G).

LDs are organelles with major functions in inflammatory mediator production and innate signaling in immune cells. To evaluate if LDs contribute to SARS-CoV-2-induced inflammation, monocytes were pre-treated with A922500, the secreted levels of lipid mediators, cytokine and chemokines were measured 24 h after infection. It has been well established that LDs are organelles that compartmentalize the eicosanoid synthesis machinery and are sites for eicosanoid formation (Bozza et al., 2011). Here, we demonstrated that SARS-CoV-2 infection increased LTB_4_ and cysLT production in comparison with uninfected monocytes (Fig 3D). The pretreatment with DGAT-1 inhibitor A922500 reduced the synthesis of both lipid mediators by infected cells (Fig. 3D). These data point out the importance of LD for the production of these pro-inflammatory lipid mediators. We also observed that A922500 treatment downregulated the chemokines IL-8 and CXCL10, and the pro-inflammatory cytokines IL-6, TNFα and IL-12 (Fig. 3E and F), without affecting the anti-inflammatory cytokine IL-10 (Fig. 3G). In addition to lowering the pro-inflammatory mediators, inhibition of LDs may shift the inflammatory profile by increasing the anti-inflammatory cytokine IL-4 (Fig. 3G).

Altogether, our data indicate that LDs have important functions in the modulation of inflammatory mediators production in SARS-CoV-2-infected monocytes and suggest that LD inhibition may reduce the exaggerated inflammatory process caused by the cytokine storm.

### Lipid droplets are sites for SARS-CoV-2 replication

The up regulation of the lipid metabolism and LD biogenesis by the new SARS-CoV-2 suggest that the virus may explore host metabolism to favor it is replication using the LDs as a replication platform, as demonstrated for HCV (Boulant et al., 2007; Camus et al., 2013; Lee et al., 2019) and DENV (Samsa et al., 2009). To evaluate this, we used a VERO E6 cell line that has a highly replicative capacity.

For these experiments, we pre-treated the VERO cells with a range of concentrations of DGAT-1 inhibitor A922500 (0.1 - 50 μM) for 2 hours, followed by infection with SARS-CoV-2 (MOI 0.01) for 24 hours. The supernatant was used to perform a plaque assay. Here, we observed that A922500 significantly inhibited SARS-CoV-2 replication in a dose dependent manner with an IC50 of 3.78 μM (Fig. 4A and B).

**Fig 4.**
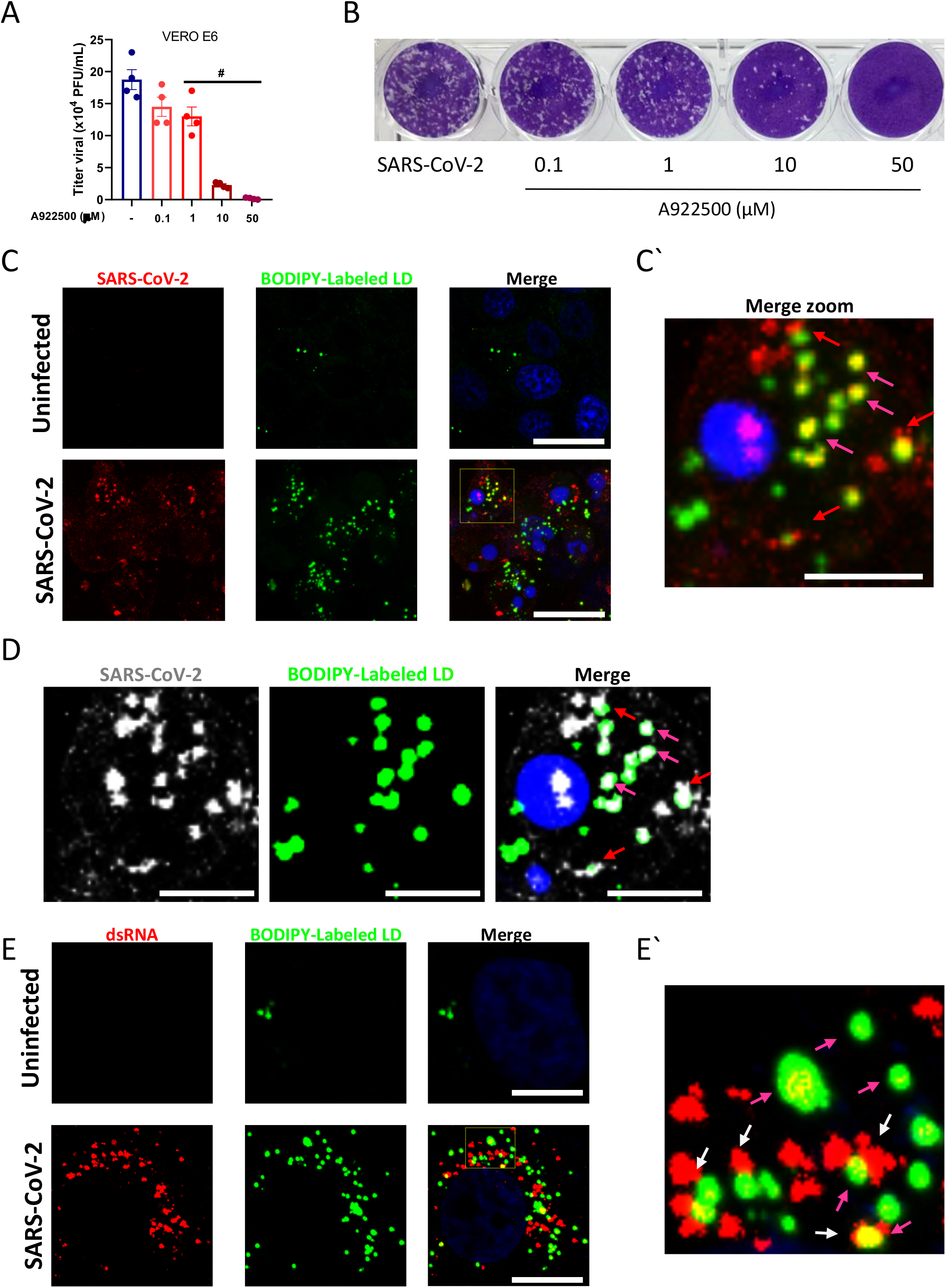
Lipid droplets is necessary for SARS-CoV-2 replication in VERO E6. VERO E6 were pre-treated with DGAT-1 inhibitor A922500 with different concentrations (0.1, 1, 10 and 50μM) for 2 hours before the infection with SARS-CoV-2 with MOI of 0.01 for 24h. (A) Viral replication was determinate by Plaque assay. (B) Representative Plaque assay. (C-E) Immunofluorescence analyses of VERO E6 after SARS-CoV-2 infection with MOI of 0.01 for 48h. (C) The virus was detected by indirect immunofluorescence using convalescent donor serum (Red or white) or (E) the double strain RNA was detected by indirect immunofluorescence by J2 antibody (Red), the lipid droplets were stained with BODIPY 493/503 (Green) and nuclei stained with DAPI (Blue). (C’ and E’) Representative zoom images. Data are expressed of four independent experiments for SARS-CoV-2 replication and three for immunofluorescent analyse. #p <0.05 versus A922500 treated cells. Scale bar 20μm.

To gain insights on the interaction of the SARS-CoV-2 with LDs we labeled the virus using immune serum from a convalescent COVID-19 patient that exhibit high anti-SARS-CoV-2 titers. For that, we stained the LDs using a BODIPY probe and analyzed the co-localization between the viral proteins and LDs by confocal microscopy. As shown in figure 4C, intense immunoreactivity (red) was obtained in SARS-CoV-2 infected cells, whereas no labeling was observed in uninfected cells, indicative of specific SARS-CoV-2 labeling with COVID-19 convalescent serum (Fig. 4C). As observed for monocytes and lung cells, Vero E6 infected cells increased LD biogenesis (green). Then, we examined the spatial relationship between SARS-CoV-2 and LDs. Confocal analysis showed a close apposition of SARS-CoV-2 immunoreactivity with BODIPY-labeled LDs (red arrows) and also co-localization of viral protein(s) with BODIPY-labeled LDs (yellow; fuchsia arrows) in the infected cells (Fig. 4C-D).

Accumulating evidence indicate that host LDs play an important role in virus replicative cycle, including as hubs for viral genome replication and viral particle assembling (Laufman et al., 2019; Lee et al., 2019; Miyanari et al., 2007; Samsa et al., 2009). To assess if LDs are associated with SARS-CoV-2 replication, we use a specific antibody for double stranded (ds)-RNA (J2 clone). As shown in figure 4E, we observed strong labeling of the ds-RNA in cells infected with the SARS-CoV-2 compared to uninfected cells. Similar to labeling detected with convalescent COVID-19 polyclonal serum, we observed close apposition and/or co-localization between BODIPY-labeled LDs and ds-RNA (Fig. 4E and E’).

Collectively, our data suggest that SARS-CoV-2 uses LDs as a replication platform, and establish that pharmacological targeting of LD formation inhibit SARS-CoV-2 replication, emerging as a potential strategy for antiviral development.

## Discussion

Most positive-strand RNA virus are able to modulate the host lipid metabolism and to highjack LDs to enhance their fitness and replication/particle assembling capacity (Herker and Ott, 2012; Pereira-Dutra et al., 2019). The pathways and mechanisms used may vary according to the virus and the host cell infected. The mechanisms and pathways explored by SARS-CoV-2 to support its replication within host cells are still largely unknown. Here we provide evidence that LDs participate at two levels of host pathogen interaction in SARS-CoV-2 infection: first, they are important players for virus replication; and second, they are central cell organelles in the amplification of inflammatory mediator production. First, we demonstrated that SARS-CoV-2 modulates pathways of lipid uptake and lipogenesis leading to increased LD accumulation in human host cells. We further showed that LDs are in close proximity with SARS-CoV-2 proteins and replicating genome, a finding suggestive that LDs are recruited as part of replication compartment. Second, we showed that inhibition of DGAT-1 blocked LD biogenesis, and reduced virus replication, cell-death and pro-inflammatory mediator production.

LD biogenesis is a multi-mediated and highly coordinated cellular process that requires new lipid synthesis and/or lipid uptake and remodeling, but the molecular mechanisms involved in LD formation during inflammation and infection are still not completely understood. Here, we showed the increased expression of SREBP-1 and the nuclear receptor PPARγ after SARS-CoV-2 infection indicative of reprogramming of cells towards a lipogenic phenotype. Accordingly, increased expression of SREBP-1 has been reported after respiratory viruses including MERS-CoV, SARS-CoV, and shown to regulate the increase of the LD and the accumulation of the cholesterol during the infection (Yuan et al., 2019). Consistently, targeting the SREBP-associated lipid biosynthetic pathways were shown to have antiviral properties (Yuan et al., 2019). The transcription factor PPARγ is activated by lipid ligands and promotes the expression of proteins involved in lipid homeostasis and LD biogenesis, and has been implicated in infectious and non-infectious LD biogenesis in monocytes/macrophages (Almeida et al., 2014; Souza-Moreira et al., 2019). Based on these data we can suggest that these two transcription factors are critical for SARS-CoV-2 infection, favoring the lipid synthesis and LD formation. One important gene up regulated by PPARγ is the membrane receptor CD36 (Cheng et al., 2016). CD36 plays an important role in the transport and uptake of long-chain fatty acids into cells and participates in pathological processes, such as metabolic disorders and infections (Febbraio et al., 2001). Previous reports showed that CD36 levels are increased in HCV and HIV-1 infections (Berre et al., 2013; Meroni et al., 2005) and that it facilitates the viral attachment on host cell membrane contributing to viral replication (Cheng et al., 2016). Our results demonstrated that SARS-CoV-2 infection increase the CD36 expression in monocytes, suggesting the increase of lipids uptake can contribute to LD formation, observed after the infection.

Numerous studies established LDs as key organelles during +RNA viruses replicative cycle (Herker and Ott, 2012). Here, we observed strong labeling of the SARS-CoV-2 proteins and ds-RNA intimately associated to the LD and in some cases colocalizing with LD. This fact highly suggests that SARS-CoV-2 recruits LDs to replication compartments and could use them as building blocks to fuel its own replication. Indeed, recent studies have shed light on active mechanisms of LD recruitment to viral replication compartments with bi-directional content exchange and essential functions to replication and virus particle assembly (Laufman et al., 2019; Lee et al., 2019).

DGAT-1, the key enzyme for triacylglycerol synthesis, is critical for LD biogenesis and mediate viral protein trafficking to LD by HCV and other viruses. Moreover, pharmacological suppression of DGAT1 activity inhibits HCV replication at the assembly step (Camus et al., 2013; Herker et al., 2010). We observed that DGAT-1 expression increases after SARS-CoV-2 infection and that this enzyme can contribute for the LD remodeling in the host cells. Pharmacological inhibitors of lipid metabolism protein are able to modulate the LD formation. Therefore, we used the DGAT-1 inhibitor (A22500) during SARS-CoV-2 infection and observed that this treatment reduced the LD biogenesis in monocytes and A549 cells, as well as decrease the viral load of SARS-CoV-2 in monocytes. Importantly, pharmacologically suppressing DGAT1 activity dose dependently inhibited SARS-CoV-2 infectious particle formation in VERO E6 cells with an IC50 of 3.78 μM. Thus, suggesting that DGAT-1 activity and LD formation are crucial to SARS-CoV-2 replication and assembly in these cells.

Dysregulated monocyte responses are pivotal in the uncontrolled production of cytokines during the infection with respiratory viruses, such as influenza A virus (Gao et al., 2013; Peschke et al., 1993). Dysregulated immune response with key involvement of monocytes, and increased pro-inflammatory cytokine/chemokine production is also observed during severe COVID-19 and is associated with the outcome of the disease (Coperchini et al., 2020; Zhou et al., 2020a). SARS-CoV-2 infection of human monocytes in vitro recapitulate most of the pattern of inflammatory mediator production associated with COVID-19 severity, including the enhancement of the IL-6 and TNFα levels, and the consistent cell death, measured by LDH release (Fintelman-Rodrigues et al., 2020; Temerozo et al., 2020; Zhou et al., 2020a). We showed that SARS-CoV-2 infection generated a large amount of inflammatory lipid mediators, and cytokine synthesis by monocytes. Blockage of DGAT-1 activity lead to inhibition of the LDs and significantly reduced leukotriene production and pro-inflammatory cytokines released by monocytes, suggesting an important role for LDs to control the inflammatory process, and consequently to prevent the cell death-related with the uncontrolled inflammation. This finding is in agreement with the well-established role of LDs in inflammation and innate immunity (Bozza and Viola, 2010; Pereira-Dutra et al., 2019). Therefore, our data support a role for LD in the heightened inflammatory production triggered by SARS-CoV-2 and conversely, inhibition of LD biogenesis by targeting DGAT1 activity may have beneficial effects on disease pathogenesis.

In summary, our data demonstrate that SARS-CoV-2 triggers reprograming of lipid metabolism in monocytes and other cells leading to accumulation of LDs favoring virus replication. The inhibition of LD biogenesis modulates the viral replication and the pro-inflammatory mediator production. Therefore, our data support the hypothesis that SARS-CoV-2 infection increases the expression of the lipid metabolism-related proteins for their own benefit towards replication and fitness. Although, further studies are certainly necessary to better characterize the full mechanisms and importance of the LDs during the SARS-CoV-2 infection, our findings support major roles for LDs in SARS-CoV-2 replicative cycle and immune response. Moreover, the finding that the host lipid metabolism and LDs are required for SARS-CoV-2 replication suggests a potential strategy to interfere with SARS-CoV-2 replication by blocking the DGAT1 and other lipid metabolic pathway enzymes.

## Acknowledgments

We thank the Hemotherapy Service from Hospital Clementino Fraga Filho (Federal University of Rio de Janeiro, Brazil) for providing buffy-coats. The authors thank the confocal imaging facility from the Rede de Plataformas Tecnológicas FIOCRUZ and Dr. Carmen Beatriz Wagner Giacoia Gripp for assessments related to BSL3 facility. This work was supported by grants from Inova program Fiocruz, Fundação de Amparo à Pesquisa do Estado do Rio de Janeiro (FAPERJ), Conselho Nacional de Desenvolvimento Científico e Tecnológico (CNPq) and Coordenação de Aperfeiçoamento de Pessoal de Nível Superior (CAPES) granted for Patrícia T. Bozza, Thiago Moreno L. Souza, Dumith Chequer Bou-Habib and Fernando A. Bozza.

## Author Contribution

Conceived the study: SSGD, VCS, FAB, TMLS, PTB; Designed the experiments: SSGD, VCS, TMLS; PTB; Performed the experiments: SSGD, VCS, ACF, CQS, NFR, JRT, LT, EB, MM, CSF, IGAQ, PPM, EH, CRRP; Analyzed the data: SSGD, VCS, DCBH, TMLS, PTB; Wrote the paper: SSGD, VCS, TMLS, PTB. All authors reviewed and approved the manuscript.

The authors declare no competing financial interests.

## Methodology

### Cells, virus and reagents

Blood were obtained from RT-PCR-confirmed COVID-19 patients and SARS-CoV-2-negative health volunteers. Human monocytes were isolated from peripheral blood mononuclear cells (PBMCs) using density gradient centrifugation (Ficoll-Paque, GE Healthcare). The PBMC were resuspended in PBS containing 1 mM EDTA and 2 % fetal bovine serum (FBS; GIBCO) to the concentration of 10^8^ cells/mL. The cells were incubated with anti-CD14 antibodies (1:10) for 10 min and magnetic beads-conjugates (1:20) for additional 10 min, followed by magnetic recovery of monocytes for 5 min. Recovered monocytes were resuspended in PBS containing 1 mM EDTA and 2 % FBS and subjected to two more rounds of selection in the magnet according to the manufacturer’s instructions (Human CD14+ selection kit, Easy Sep; StemCell). The purity of monocyte preparations (>98% CD14+ cells) was confirmed through flow cytometry.

Human primary monocyte was obtained through plastic adherence of PBMCs. Briefly, PBMCs were isolated by Ficoll-Paque from peripheral blood or from buff-coat preparations of healthy donors. PBMCs (2 × 10^6^) were plated onto 48-well plates in low glucose Dulbecco’s modified Eagle’s medium (DMEM; GIBCO). After 2 hours of the plaque, non-adherent cells were washed out and the remaining monocytes were maintained for 24 hours in DMEM containing 5% inactivated male human AB serum (HS; Merck) and 100 U/mL penicillin-streptomycin (P/S; GIBCO) at 37 °C in 5 % CO_2_. The purity of human monocytes was above 90 %, as analyzed by flow cytometry analysis (FACScan; Becton Dickinson) using anti-CD3 (BD Biosciences) and anti-CD16 (Southern Biotech) monoclonal antibodies.

Human lung epithelial carcinoma cell line (A549 - ATCC/CCL-185) and African green monkey kidney (Vero subtype E6) were cultured in high glucose DMEM supplemented with 10% FBS and 100 U/mL P/S, and were incubated at 37 °C in 5 % CO_2_.

Human lung microvascular endothelial cell line (HMVEC-L - LONZA/CC-2527) was maintained following the manufacturer’s instructions. The cells were cultured in endothelial growth medium (EGM™-2MV BulletKit™, Clonetics) supplemented with 5 % fetal bovine serum (FBS, Clonetics) and cells were incubated at 37 °C and 5 % CO_2_.

SARS-CoV-2 was originally isolated from nasopharyngeal swabs of confirmed case from Rio de Janeiro/Brazil (GenBank accession no. MT710714). The virus was amplified in Vero E6 cells in high glucose DMEM supplemented with 2% FBS, incubated at 37°C in 5% CO_2_ during 2 to 4 days of infection. Virus titers were performed by the tissue culture infectious dose at 50% (TCID_50_/mL) and the virus stocks kept in −80 °C freezers. According to WHO guidelines, all procedures involving virus culture were performed in biosafety level 3 (BSL3) multiuser facility.

### Infections and virus titration

After 24h of cell plating, the SARS-CoV-2 infections were performed at MOI of 0.01 in all cells with or without pre-treatment with the pharmacological inhibitor of DGAT-1 (A922500 – Sigma CAS 959122-11-3) for two hours. The Plaque-forming Assay was performed for virus titration in VERO E6 cells seeded in 24-well plates. Cell monolayers were infected with different dilutions of the supernatant containing the virus for 1h at 37°C. The cells were overlaid with high glucose DMEM containing 2% FBS and 2.4% carboxymethylcellulose. After 3 days, the cells were fixed with 10% formaldehyde in PBS for 3h. The cell monolayers were stained with 0.04% crystal violet in 20% ethanol for 1h. The viral titer was calculated from the count of plaques formed in the wells corresponding to each dilution and expressed as plaque forming unit per mL (PFU/mL).

### Lipid droplet staining

Human primary monocytes, A549 cell line, and HMVEC cell line were seeded in coverslips. The cells infected or not were fixed using 3.7% formaldehyde. In addition, after isolation, the monocytes from COVID-19 patients were fixed using 3.7% formaldehyde and adhered in coverslips through cytospin (500 x g for 5 min). The LDs were stained with 0.3% Oil Red O (diluted in 60% isopropanol) for 2 min at room temperature. The coverslips were mounted in slides using an antifade mounting medium (VECTASHIELD®). Nuclear recognition was based on DAPI staining (1 μg/mL) for 5 min. Fluorescence was analyzed by fluorescence microscopy with an 100x objective lens (Olympus, Tokyo, Japan). The numbers of LDs were automatically quantified by ImageJ software analysis from 15 aleatory fields.

### Immunofluorescence staining

VERO E6 cells were seeded in coverslips and after 48h were fixed using 3.7% formaldehyde. Cells were rinsed three times with PBS containing 0.1 M CaCl2 and 1 M MgCl2 (PBS/CM) and then permeabilized with 0.1% Triton X-100 plus 0.2% BSA in PBS/CM for 10 min (PBS/CM/TB). Cells were stained with convalescent serum from a patient to identify with COVID-19 at 1:500 dilution for overnight, followed by a human anti-IgG-Alexa 546 at 1:1000 dilution for 1 h. The double-RNA was labeling by mouse monoclonal antibody J2 clone - Scicons (Schönborn et al., 1991) at 1:500 dilution for overnight, followed by a mouse anti-IgG-Dylight 550 at 1:1000 dilution for 1h. LDs were stained with BODIPY493/503 dye (dilution 1:5000 in water) for 5 min. The coverslips were mounted in slides using an antifade mounting medium (VECTASHIELD®). Nuclear recognition was based on DAPI staining (1 μg/mL) for 5 min. Fluorescence was analyzed by fluorescence microscopy with an 100x objective lens (Olympus, Tokyo, Japan) or Confocal Microscopy (Laser scanning microscopy LSM710 Meta, Zeiss).

### SDS-PAGE and Western blot

After 24h of SARS-CoV-2 infection, monocytes were harvested using ice-cold lysis buffer (1% Triton X-100, 2% SDS, 150 mM NaCl, 10 mM HEPES, 2 mM EDTA containing protease inhibitor cocktail - Roche). Cell lysates were heated at 100 °C for 5 min in the presence of Laemmli buffer (20% β-mercaptoethanol; 370 mM Tris base; 160 μM bromophenol blue; 6% glycerol; 16% SDS; pH 6.8). Twenty μg of protein/sample were resolved by electrophoresis on SDS-containing 10% polyacrylamide gel (SDS-PAGE). After electrophoresis, the separated proteins were transferred to nitrocellulose membranes and incubated in blocking buffer (5% nonfat milk, 50 mM Tris-HCl, 150 mM NaCl, and 0.1% Tween 20). Membranes were probed overnight with the following antibodies: anti-PPARγ (Santa Cruz Biotechnology, #SC-7196 - H100), anti-CD36 (Proteintech-18836-1-AP), anti-SREBP-1 (Ab-28481), anti-DGAT-1 (Santa Cruz Biotechnology, #SC-271934) and anti-β-actin (Sigma, #A1978). After the washing steps, they were incubated with IRDye - LICOR or HRP-conjugated secondary antibodies. All antibodies were diluted in blocking buffer. The detections were performed by Supersignal Chemiluminescence (GE Healthcare) or by fluorescence imaging using the Odyssey system. The densitometries were analyzed using the Image Studio Lite Ver 5.2 software.

### Measurement of viral RNA load

Supernatants from monocytes were harvested after 24h of SARS-CoV-2 infection and the viral RNA quantified through RT-PCR. According to manufacter’s protocols, the total RNA from each sample was extracted using QIAamp Viral RNA (Qiagen®). Quantitative RT-PCR was performed using QuantiTect Probe RT-PCR Kit (Quiagen®) in a StepOne™ Real-Time PCR System (Thermo Fisher Scientific).

Amplifications were carried out containing 2× reaction mix buffer, 50 μM of each primer, 10 μM of probe, and 5 μL of RNA template in 15 μL reaction mixtures. Primers, probes, and cycling conditions recommended by the Centers for Disease Control and Prevention (CDC) protocol were used to detect the SARS-CoV-2 (CDC, 2020). For virus quantification it was employed the standard curve method. Cells of each sample were counted before the PCR analyses for normalization. The Ct values for this target were compared to those obtained to different cell amounts, 10^7^ to 10^2^, for calibration.

### Measurements of inflammatory mediators and LDH activity

The monocyte supernatant was obtained after 24 hours of SARS-CoV-2 infection with or without pre-treatment with A922500 (10 μM). Cytokines and chemokines were measured in the supernatant by ELISA following the manufacturer’s instructions (Duo set, R&D). LTB_4_ and cysLT were measured in the supernatant by EIA following the manufacturer’s instructions (Cayman Chemicals). Cell death was determined according to the activity of lactate dehydrogenase (LDH) in the culture supernatants using a CytoTox® Kit according to the manufacturer’s instructions (Promega, USA).

### Ethics statement

Experimental procedures involving human cells from healthy donors were performed with samples obtained after written informed consent and were approved by the Institutional Review Board (IRB) of the Oswaldo Cruz Foundation/Fiocruz (Rio de Janeiro, RJ, Brazil) under the number 397-07. Experimental procedures involving human patient cells were performed with samples obtained after written informed consent from all participants or patients’ representatives according to the study protocol approved by the National Review Board (CONEP 30650420.4.1001.0008).

### Statistical analysis

Data are expressed as mean ± standard error of the mean (SEM) at least of three and maximum of five independent healthy donors. The paired two-tailed *t*-test was used to evaluate the significance of the two groups. Multiple comparisons among three or more groups were performed by one-way ANOVA followed by Tukey’s multiple comparison test. p values < 0.05 were considered statistically significant when compared SARS-CoV-2 infection to the uninfected control (*) group or SARS-CoV-2 infection with A922500 treat group (#).

**Fig S1.**
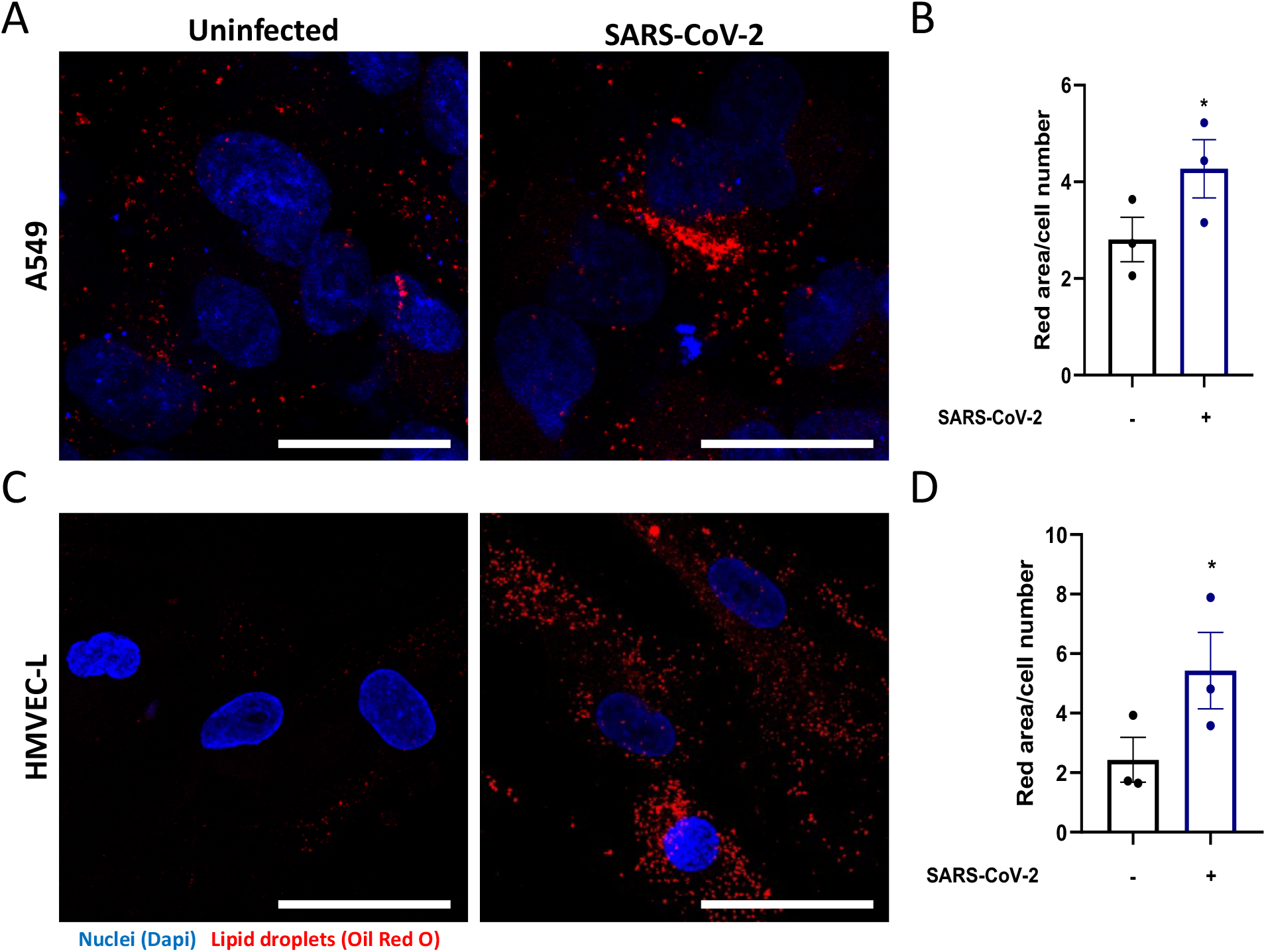
SARS-CoV-2 induces an increase of the LD biogenesis in different human pulmonary cell lines. Human pulmonary cell lines were infected with SARS-CoV-2 at MOI of 0.01 for 48h. (A and C) LDs were captured by fluorescent microscopy after Oil Red O staining (Red) and nuclei stained with DAPI (Blue). (B and D) LDs were evaluated by ImageJ software analysis by the measurement of the fluorescent area. Data are expressed of three independent experiments. *p <0.05 versus uninfected cells. Scale bar 20μm.

